## Supplementary_Methods&Figures for "Microglia-neuron communication at nodes of Ranvier depends on neuronal activity through potassium release and contributes to myelin repair"

#### Tissue preparation

*Ex vivo cerebellar slices*

Cultured slices were fixed while attached to the membrane with either 4% or 1% paraformaldehyde (PFA) for 30 min at room temperature (RT) and washed with PBS.

*In vivo mouse brain and spinal cord*

For spinal cord preparation, the tissue was dissected following perfusion and post-fixed in PFA 2% for 30min, washed in PBS and incubated in PBS, 15% sucrose for 3 days at 4°C. Spinal cords were then embedded in PBS, 15% Sucrose, 4% gelatin and frozen in isopropanol onto carbonic ice.

For adult as well as P12 myelinating mouse brain tissue, animals were perfused with PFA 2% and the brain was dissected and post-fixed in PFA 2% for 30min, washed in PBS and incubated in PBS with successively 7, 15 and 30% sucrose for 3 days at 4°C for cryoprotection. The tissues were then embedded in O.C.T (Tissue-Tek, Sakura).

#### Tissue sectioning

Mouse spinal cords and brains were cut longitudinally and sagitally respectively. Human tissues (healthy donors, UK MS Society Tissue Bank at Imperial College, London, under ethical approval by the National Research Ethics Committee 08/MRE09/31) were obtained as snap frozen blocks. Sections were cut using a cryostat Leica CM 1950 (30 and 15 µm thick for mouse and human respectively), collected on Superfrost+ slides and stored at -80°C until used.

#### Immunhistostainings

*Immunofluorescent stainings*

Human sections were first fixed in 4% PFA for 5 minutes. For myelin protein staining, tissues were preincubated in absolute ethanol at -20°C for 20 min. Tissues were blocked in PBS, 5-10% Normal Goat Serum (50-062Z; Thermo Fisher Scientific), 0,2-0,4% Triton X-100 (RDTH), then incubated with primary antibodies in blocking solution overnight at room temperature, washed in PBS and incubated with secondary antibodies for 1-3 hours at RT in the dark. When applicable, tissues were immersed for 30s in Hoechst solution (10 µg/ml,

Euromedex). They were finally mounted under coverslip (VWR) with Fluoromount (Southern Biotech).

***Chromogenic immunohistochemistry on post-mortem brain tissues***

Snap frozen tissue blocks were rehydrated with PBS. Sections were then immersed in 4% PFA for 10 minutes before elimination of endogenous peroxidase activity with 0.1% H2O2 (Sigma Aldrich) in PBS for 20 minutes. Blocking was performed using 5% Normal Goat Serum before incubation with primary antibodies diluted in PBS containing 0.05% Triton 100-X and 5% sera. The biotinylated secondary antibody targeting one of the primary antibodies (Vector Laboratories) was visualized with the avidin-biotin horseradish peroxidase complex (Dako, Biotin Blocking System) followed by 3,3'-diamino-benzidine (DAB) (Vector Laboratories) as substrate. The second primary antibody was detected with the ABC-alkaline phosphatase detection system (Vector Laboratories), using Vector Blue as the substrate. Images were captured with a QICAM digital camera (QImaging Inc.).

**Imaging**

*Confocal microscopy for ex vivo cultured slices*

Confocal microscopy was performed using an FV-1200 Upright Confocal Microscope and a Leica inverted SP8 with 63x oil immersion objectives with 1.40 numerical aperture, using respectively metamorph and LasX software. For each acquisition, stacks of 1024x1024 pixel images (160.6 µm x 160.6 µm), including at least 10 Z-series with a z-step of 0.30 µm, were acquired using 405, 488, 552 and 638 laser lines or a white laser with optimized wavelengths at the peak of excitation for each fluorophore.

*Confocal microscopy for in vivo tissue*

Confocal microscopy was performed using an inverted Leica SP8 with 40x or 63x oil immersion objectives with 1.30 and 1.40 numerical aperture respectively, using LasX software. For quantification, 1024x1024 pixel images were acquired using a 40X oil-objective with a numerical zoom of 2, corresponding to a 211.0 µm x 211.0 µm final area, and 10 sections were acquired with a step of 0.30 µm. For 3D reconstruction, images were acquired using the 63x oil immersion objective, and a z-step of 0.20 µm. Deconvolution was carried out using Huygens software (v.17.10). Following deconvolution, the surface of each structure was reconstructed in 3D using Imaris software (GraphPad, Bitplane, v.9.2).

Figures were made using Photoshop (Adobe, version CC).

**Electrophysiology**

Myelinated cultured slices (10-13 DIV) were transferred to a recording chamber and continuously superfused with oxygenated (95% $O_2$ and 5% $CO_2$) aCSF containing (in mM): 124 NaCl, 3 KCl, 1.25 $NaH_2PO_4$, 26 $NaHCO_3$, 1.3 $MgSO_4$, 2.5 $CaCl_2$, and 15 glucose (pH 7.4). Purkinje cells were visualized under differential interference contrast optics using a 63X N.A 1 water immersion objective. Loose patch voltage clamp recordings of the spontaneous firing activity was performed at 32-34°C with a borosilicate glass pipette filled with aCSF. Signals were amplified with a Multiclamp 700B amplifier (Molecular devices), sampled and filtered at 10 kHz with a Digidata 1550B (Molecular Devices). Data were acquired with the pClamp software (Molecular devices, version 10.4). To avoid any alteration of the spontaneous firing frequency of the cell by the patch procedure (Perkins, 2006), the holding membrane potential was set to the value at which zero current is injected by the amplifier. The resistance of the seal ($R_{seal}$) was controlled and calculated every minute from the current response to a voltage step (200 ms; -10 mV). Only recordings with a $R_{seal}$ in the range of 10 to 200MΩ and stable during the all recording procedure were included in the analysis. Spontaneous activity was tested in control condition and consecutively in the presence of apamin (500 nM) and TTX (500 nM) applied by bath perfusion. The firing rate was analysed over a two minutes recording time window using a threshold crossing spike detection in Clampfit (Molecular devices, version 10.4) and calculated as the number of action potential current divided by the duration of the recording. The instantaneous frequency was calculated as the multiplicative inverse of the interspike interval.

### Supplementary Figure and Movie legend

**Figure S1. Microglial contacts with nodes of Ranvier are less stable during demyelination.** **(A-C)** *In vivo* live-imaging from CX3CR1-GFP/Thy1-Nfasc186mCherry mouse dorsal spinal cord with an initial contact between microglia (green) and nodes of Ranvier (red, arrowheads); 1-hour movies with an acquisition every 10 minutes. **(A)** Sham animal (NaCl injection) imaging at 7 DPI (corresponding to Movie 2); **(B)** LPC-injected animal imaged at 7 DPI (demyelination, corresponding to Movie 3); **(C)** LPC-injected animal at 11 DPI (remyelination, corresponding to Movie 4). **(D)** Percentage of frames with microglia-node contact in 3-hour movies. **(E)** Longest sequence of consecutive timepoints with microglia-node contact in 3-hour movies. Each dot is a microglia-node pair. The number of microglia-node pairs imaged is indicated on each bar (n=4 to 7 animals per condition). (**A**, **B**, **C**) Scale bar 10 µm. (**D**, **E)** Kruskal-Wallis test. Bars and error bars represent the mean ± s.e.m. For detailed statistics, see Supplementary Table.

**Figure S2. Microglial cells contact nodal structures in *ex vivo* organotypic cerebellar slice culture.** **(A, B)** Microglia contact nodal structures during myelination in CX3CR1$^{GFP/+}$ mouse cerebellum *in vivo* (**A**, P12) and *ex vivo* (**B**, 4 DIV). Both mature nodes of Ranvier (filled arrowheads) and immature nodal structures (node-like clusters and heminodes, empty arrowheads) are contacted. **(C)** Percentage of nodal structures contacted by microglia in myelinating slices *ex vivo* (n=4 animals). **(D)** Percentage of mature and immature nodal structures contacted in myelinating condition *ex vivo* (n=4 animals per condition). **(E, F)** Microglia also contact nodal structures in myelinated **(E)** and remyelinating slices **(F)** *ex vivo* (11 DIV). Arrowheads indicate the nodes of Ranvier contacted by microglia. **(G)** Percentage of nodal structures contacted by microglia in myelinated (ctrl) vs remyelinating (rem) slices, (n=6 animals per condition). **(H)** Percentage of mature and immature nodal structures contacted in remyelinating condition (n=6 animals per condition). Scale bars: **(A, B)** 5µm, **(E, F, I, J)** 10 µm. **(D)** Wilcoxon matched pairs test; **(G, H)** Paired t-test. $^{*}P < 0.05$, $^{**}P < 0.01$, $^{***}P < 0.001$, $^{****}P < 0.0001$, ns: not significant; bars and error bars represent the mean ± s.e.m. For detailed statistics, see Supplementary Table.

**Figure S3. Live-imaging of microglia-node contact in organotypic cerebellar slice culture.** (A) Live images of a microglial cell (GFP$^{+}$) at two different timepoints, and the corresponding overlay showing the dynamics of the microglial cell, with extending (pink) and retracting (cyan) processes (arrowheads: most dynamic processes). **(B)** Myelinated CX3CR1$^{GFP/+}$ cerebellar slice with a Purkinje cell expressing β1Na$_V$mCherry following lentiviral transduction. (i) Live-image showing a node (mCherry$^+$, filled arrowhead) contacted by a microglial cell, and a non-contacted node (empty arrowhead). (ii) Summed projection of the movie showing the axon trajectory. **(C)** Maximum duration without contact between a microglial process and an internode or a node in myelinated slices (10 minutes acquisition, internode: n=32 contacts from 16 animals, node: n=9 contacts from 8 animals). **(D)** Distance between the process tip and its position at t0 for each frame, whether the process was initially contacting a node (nodal contact) or without contact (myelinated slices; wo contact: 280 measures from 14 trajectories, n=7 animals; nodal contact: 140 measures from 7 trajectories, n=7 color coded animals). **(E)** Maximum duration without contact between the tip of a microglial process and the node in myelinated vs remyelinating slices (10 minutes acquisition, ctrl: n=9 contacts from 8 animals, rem: n=6 contacts from 6 color coded animals). **(F)** Distance between the process tip and its position at t0 for each frame (remyelinating slices; wo contact: 240 measures from 12 trajectories, n=6 animals, nodal contact: 120 measures from 6 trajectories, n=6 animals). (**A**, **B)** Scale bars: 10 µm **(C, E)** Mann-Whitney test; **(D, F)** Type II Wald chi-square test. *P < 0.05, **P < 0.01, ***P < 0.001, ****P < 0.0001, ns: not significant; bars and error bars represent the mean ± s.e.m. For detailed statistics, see Supplementary Table.

**Figure S4. The microglial receptors CX3CR1, P2Y12R and P2Y13R are not required for microglia-node interaction. (A)** Microglia (GFP, in green) contact nodes (Na$_V$, in red) both in *CX3CR1*$^{GFP/+}$ and *CX3CR1*$^{GFP/GFP}$ myelinated cerebellar slices. **(B)** Percentage of nodes of Ranvier contacted by microglia in *CX3CR1*$^{GFP/+}$ vs *CX3CR1*$^{GFP/GFP}$ littermate slices (n=4 animals per condition). **(C, E)** Illustration of microglia-node contacts in myelinated slices treated with PSB0739 (**C**, P2Y12R inhibitor, 1 µM, 4-hour treatment) or MRS2211 (**E**, P2Y12R/P2Y13R inhibitor 50 µM, 4-hour treatment). **(D, F)** Percentage of nodes contacted by microglial cells in *CX3CR1*$^{GFP/+}$ myelinated slices in control vs treated condition, with PSB0739 (**D**, 1 µM) or MRS2211 (**F**, 50 µM), n=4 animals per condition. Arrowheads show the nodes of Ranvier contacted by microglial cells. Scale bars: **(A, Ciii, E)** 10 µm, **(Cvi)** 5 µm. **(B)** Mann-Whitney rank test, **(D, F)** Wilcoxon matched pairs test. *P < 0.05, **P < 0.01, ***P < 0.001, ****P < 0.0001, ns: not significant; bars and error bars represent the mean ± s.e.m. For detailed statistics, see Supplementary Table.

**Figure S5. Purkinje cells present spontaneous activity in myelinated organotypic cerebellar slices. (A)** Representative example of loose-cell attached recordings on a Purkinje cell, in control condition followed by Apamin (500 nM) and TTX (500 nM) consecutive treatments. **(B)** Instantaneous firing frequencies recorded in Purkinje cells of myelinated cerebellar organotypic slices (n=9 cells, n=7 animals, $p<0.0001$, Friedman test). **(C)** Instantaneous firing frequencies recorded in Purkinje cells in myelinated cerebellar organotypic slices after Apamin treatment, normalized by their matching control. The increased ratio of instantaneous frequencies ranges from 1.12 to 2.69 (n=9 cells, n=7 animals; center line, median; box limits, upper and lower quartiles; whiskers, min and max). *$P < 0.05$, **$P < 0.01$, ***$P < 0.001$, ****$P < 0.0001$, ns: not significant; bars and error bars represent the mean ± s.e.m. For detailed statistics, see Supplementary Table.

**Movie 1 (associated to Figure 2). Stable interaction between a node and a microglial cell along time.**
Representative example of a 3-hour movie from a CX3CR1-GFP/Thy1-Nfasc186mCherry mouse showing a stable interaction between a microglial cell (green) and a node of Ranvier (red). Acquisitions every 30 minutes. Sham animal (NaCl injection) DPI 11. Filled arrowheads indicate contact, empty arrowheads an absence of contact. Scale bar: 10µm.

**Movie 2 (associated to Figure S1). Stable interaction between a node and a microglial cell in control condition.** Representative example of a 1-hour movie from a CX3CR1-GFP/Thy1-Nfasc186mCherry mouse allowing to observe interactions between microglia in green and nodes of Ranvier in red. Acquisitions every 10 minutes. Sham animal (NaCl injection) DPI 7. Filled arrowheads indicate contact, empty arrowheads an absence of contact. Scale bar: 10µm.

**Movie 3 (associated to Figure S1). Unstable interaction between a node and a microglial cell in the perilesional tissue in demyelinating context.** Representative example of a 1-hour movie from a CX3CR1-GFP/Thy1-Nfasc186mCherry mouse allowing to observe the intermittent interaction between microglia in green and nodes of Ranvier in red. Acquisitions every 10 minutes. LPC injection DPI 7 (perilesional area, peak of demyelination). Filled arrowheads indicate contact, empty arrowheads an absence of contact. Scale bar: 10µm.

**Movie 4 (associated to Figure S1). Stable interaction between a node and a microglial cell during remyelination.** Representative example of a one-hour movie from a CX3CR1-GFP/Thy1-Nfasc186mCherry mouse allowing to observe stable interactions between microglia in green and nodes of Ranvier in red. Acquisitions every 10 minutes. LPC injection DPI 11 (remyelination). Filled arrowheads indicate contact, empty arrowheads an absence of contact. Scale bar: 10µm.

**Movie 5 (associated to Figure 4). A microglial process tip contacting an internode in a myelinated slice.** Representative example showing a microglial cell (green) initially contacting an internode (red) in a myelinated slice. Filled arrowheads indicate a contact, and empty arrowheads an absence of contact. The dashed line represents the axon. Scale bar: 5µm.

**Movie 6 (associated to Figure 4). A microglia process tip contacting a node in a myelinated slice.** Representative example showing a microglial cell (green) initially contacting a node (red) in a myelinated slice. Filled arrowheads indicate a contact, empty arrowheads an absence of contact. The dashed line represents the axon. Scale bar: 5µm.

**Movie 7 (associated to Figure 4). A microglia process tip contacting a node in a remyelinating slice.** Representative example showing a microglial cell (green) initially contacting a node (red) in a remyelinating slice. Filled arrowheads indicate a contact, empty arrowheads an absence of contact. The dashed line represents the axon. Scale bar: 5µm.

**Movie 8 (associated to Figure 6). Microglia process tip contacting a node in a myelinated slice treated with TEA.** Representative example showing a microglial cell (green) initially contacting a node (red) in a myelinated slice treated with TEA. Filled arrowheads indicate a contact, empty arrowheads an absence of contact. The dashed line represents the axon. Scale bar: 5µm.

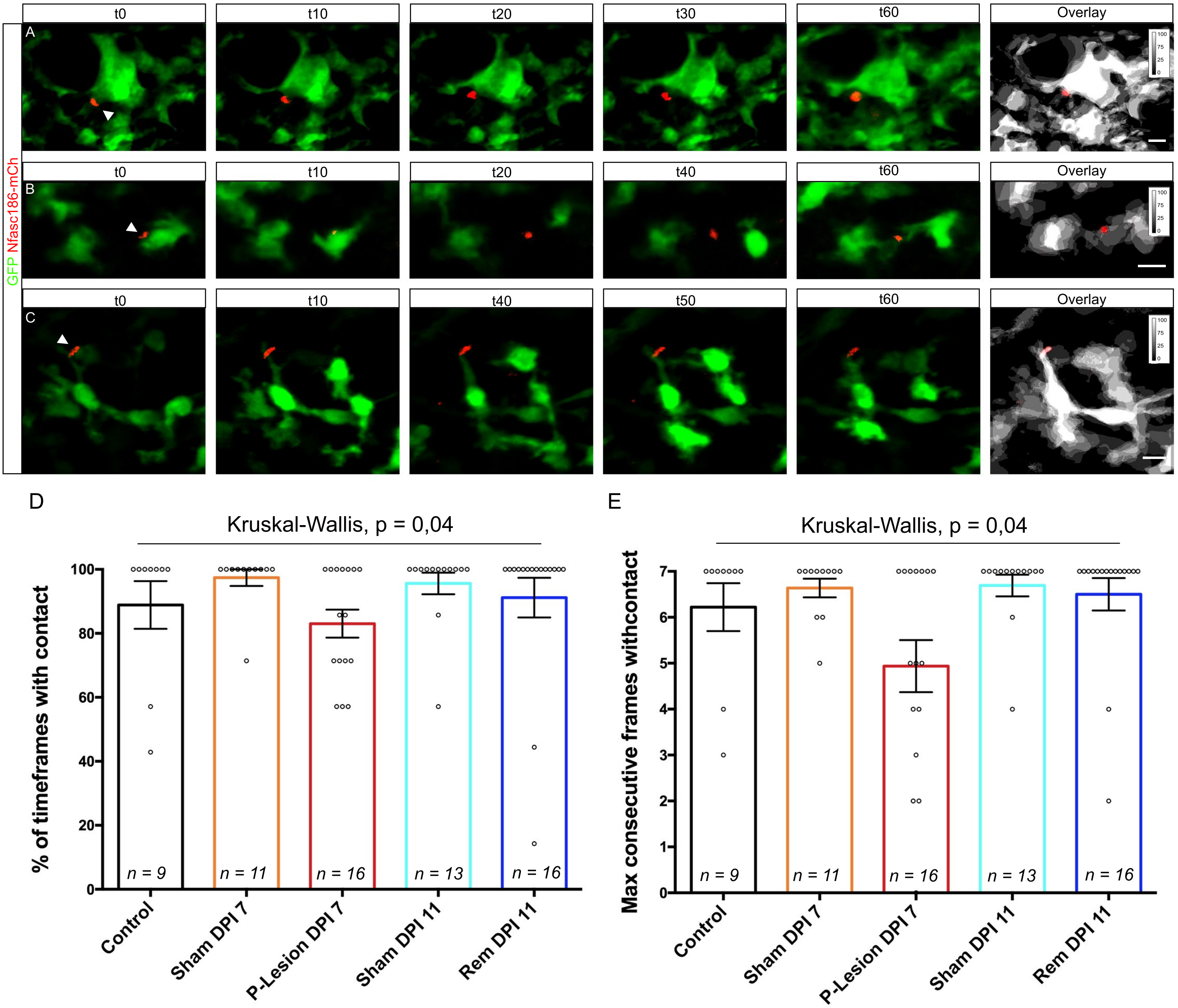

t0
t10
t20
t30
t60
Overlay
t40
t50
A
B
C
GFP Nfasc186-mCh
D
E
Kruskal-Wallis, p = 0,04
% of timeframes with contact
Max consecutive frames withcontact
n = 9
n = 11
n = 16
n = 13
n = 16
Control
Sham DPI 7
P-Lesion DPI 7
Sham DPI 11
Rem DPI 11


Figure S1

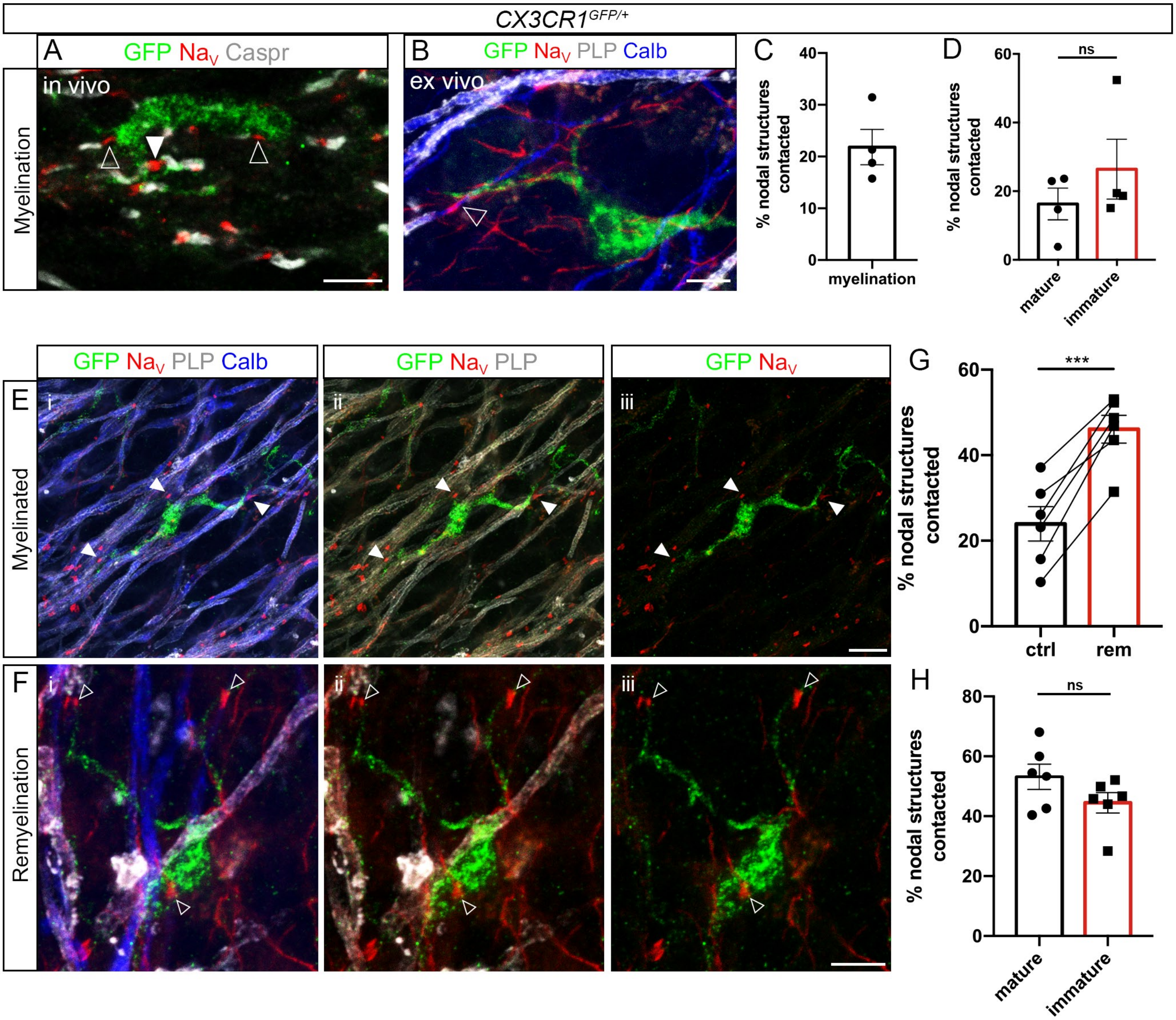

CX3CR1GFP/+
A
GFP Nav Caspr
in vivo
Myelination
B
GFP Nav PLP Calb
ex vivo
C
% nodal structures contacted
myelination
D
ns
% nodal structures contacted
mature
immature
GFP Nav PLP Calb
GFP Nav PLP
GFP Nav
E
Myelinated
i
ii
iii
G
***
% nodal structures contacted
ctrl
rem
F
Remyelination
H
ns
% nodal structures contacted
mature
immature


Figure S2

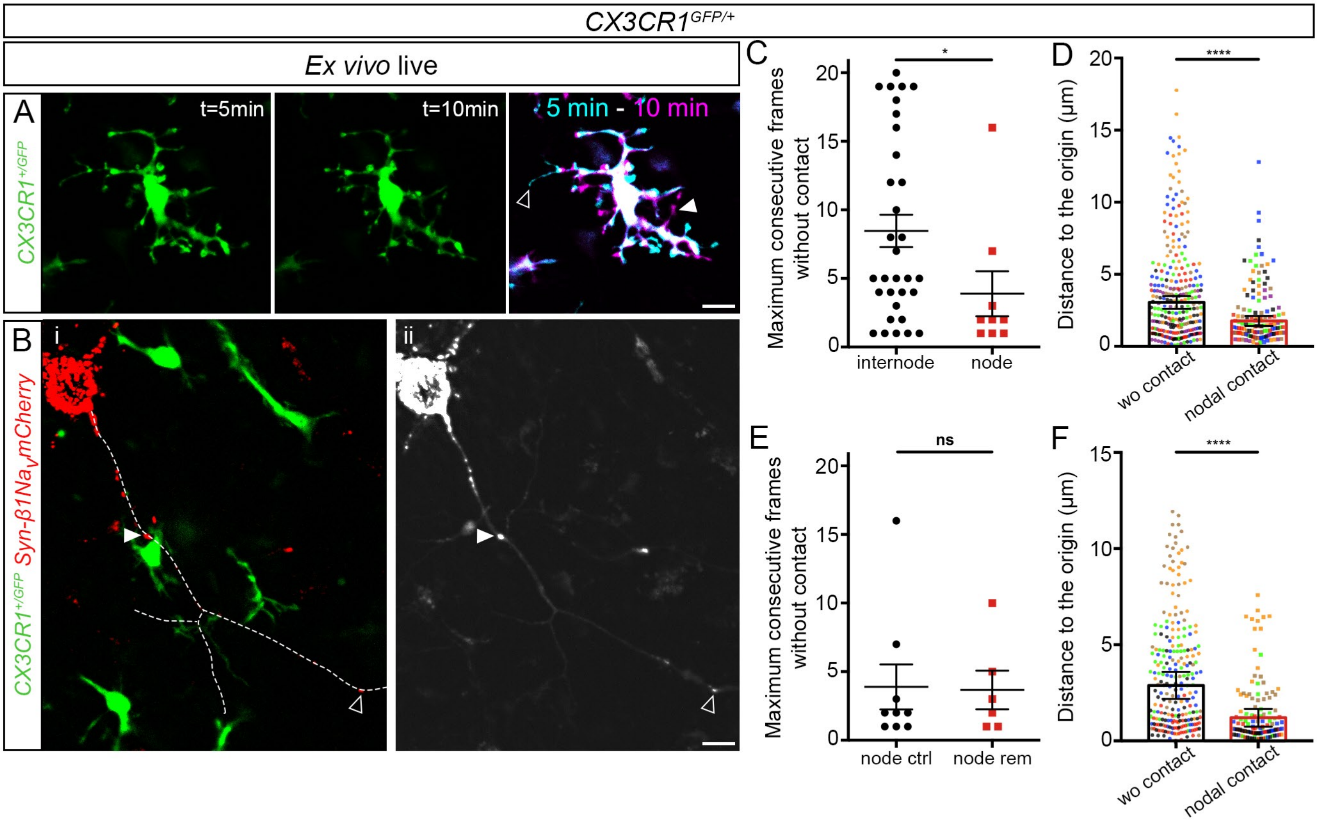
CX3CR1GFP/+
Ex vivo live
A
t=5min
t=10min
5 min - 10 min
CX3CR1+/GFP
B
i
ii
CX3CR1+/GFP Syn-β1NaVmCherry
C
Maximum consecutive frames without contact
internode
node
*
D
Distance to the origin (µm)
wo contact
nodal contact
****
E
Maximum consecutive frames without contact
node ctrl
node rem
ns
F
Distance to the origin (µm)
wo contact
nodal contact
****

Figure S3

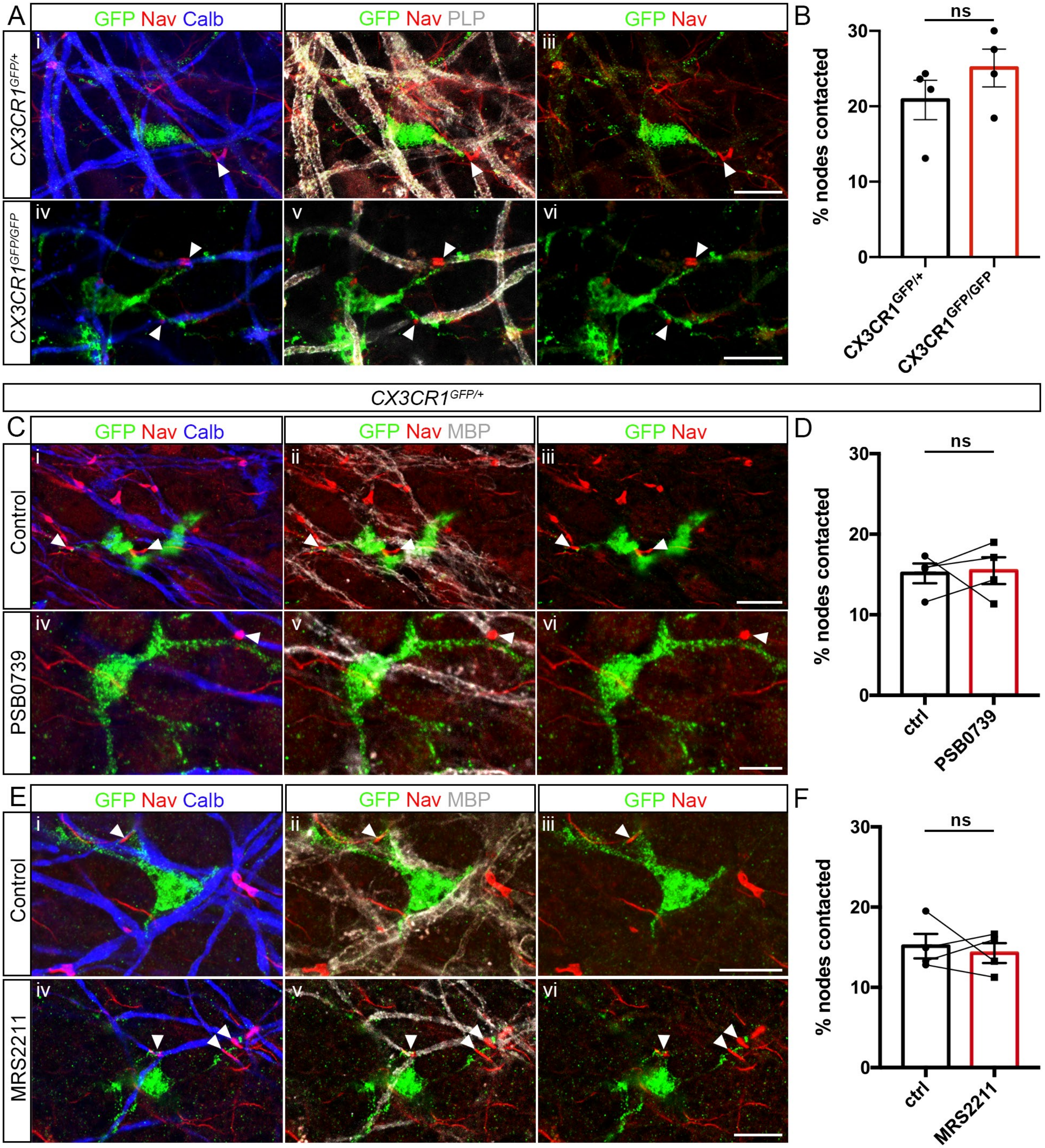
A
GFP Nav Calb
GFP Nav PLP
GFP Nav
CX3CR1GFP/+
CX3CR1GFP/GFP
i
ii
iii
iv
v
vi
B
ns
% nodes contacted
30
20
10
0
CX3CR1GFP/+
CX3CR1GFP/GFP
CX3CR1GFP/+
C
GFP Nav Calb
GFP Nav MBP
GFP Nav
Control
PSB0739
D
ns
% nodes contacted
ctrl
PSB0739
E
GFP Nav Calb
GFP Nav MBP
GFP Nav
Control
MRS2211
F
ns
% nodes contacted
ctrl
MRS2211

Figure S4

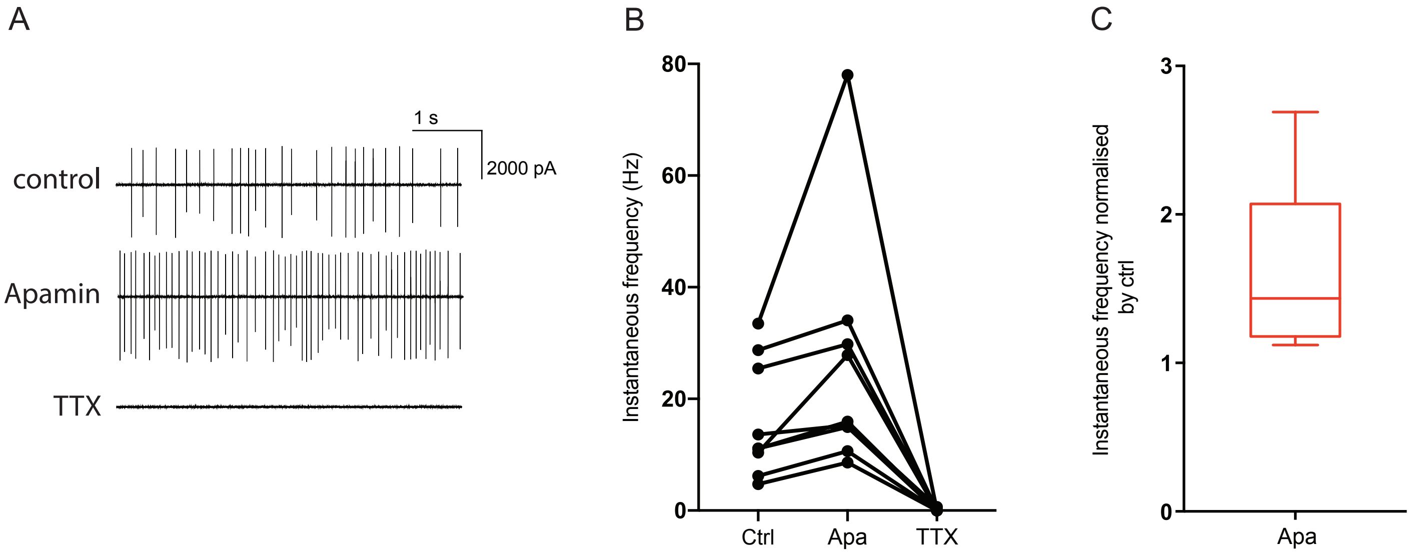
A
1 s
2000 pA
control
Apamin
TTX
B
Instantaneous frequency (Hz)
80
60
40
20
0
Ctrl
Apa
TTX
C
Instantaneous frequency normalised by ctrl
3
2
1
0
Apa

Figure S5
